# Stromal architecture directs early dissemination in pancreatic ductal adenocarcinoma

**DOI:** 10.1101/2021.02.19.431984

**Authors:** Arja Ray, Mackenzie K. Callaway, Nelson J. Rodríguez-Merced, Alexandra L. Crampton, Marjorie Carlson, Kenneth B. Emme, Ethan A. Ensminger, Alexander A. Kinne, Jonathan H. Schrope, Haley R. Rasmussen, Hong Jiang, David G. Denardo, David K. Wood, Paolo P. Provenzano

## Abstract

Pancreatic ductal adenocarcinoma (PDA) is an extremely metastatic and lethal disease. Here in both murine and human PDA we demonstrate that extracellular matrix architecture regulates cell extrusion and subsequent invasion from intact ductal structures through Tumor-Associated Collagen Signatures (TACS), resulting in early dissemination from histologically pre-malignant lesions and continual invasion from well-differentiated disease. Furthermore, we show that pancreatitis results in invasion-conducive architectures, thus priming the stroma prior to malignant disease. Analysis in novel microfluidics-derived microtissues and *in vivo* demonstrates decreased extrusion and invasion following focal adhesion kinase (FAK) inhibition, consistent with decreased metastasis. Thus, data suggest that targeting FAK or strategies to re-engineer and normalize tumor microenvironments, may have a role not only in also in very early disease but also for limiting continued dissemination from unresectable disease. Likewise, it may be beneficial to employ stroma targeting strategies to resolve precursor diseases such as pancreatitis in order to remove stromal architectures that increase risk for early dissemination.

**Impact Statement:** Collagen architectures in the tumor stroma facilitate dissemination of carcinoma cells from the earliest histologically “pre-malignant” lesions and continue to promote disease spread from well-differentiated PDA.

## Introduction

Pancreatic ductal adenocarcinoma (PDA) is a lethal disease with a dismal 5-year survival of ~9% (*1*). PDA is commonly characterized by a robust fibroinflammatory, or desmoplastic, response with hyperactivated stromal fibroblasts, robust immunosuppression, and vastly elevated extracellular matrix (ECM) deposition (*2, 3*). Apart from limiting the delivery and efficacy of chemotherapy (*4–6*) and immunotherapy (*7, 8*), the ECM of the desmoplastic stroma also plays a role in the extensive metastasis frequently observed in PDA (*9, 10*), and may influence early dissemination. Indeed, recent studies using an autochthonous mouse model of PDA demonstrate that single carcinoma cells can disseminate into the ECM-rich stroma and peripheral blood even before frank histologically-detectable malignancy (*11*). This is consistent with early disseminated cancer cells (DCCs) observed in breast cancer (*12, 13*) and decreased efficiency of dissemination in PDA following treatment with antifibrotic or anti-inflammatory agents (*7, 11*), thus challenging the concept that metastasis is always a late event in cancer progression. Hence, the desmoplastic stroma appears to be a key player in early metastatic cell dissemination, yet the mechanisms by which the fibrotic stroma aids early dissemination are unknown.

One of the main components of the dense ECM in PDA is fibrillar collagen, which is thought to be deposited largely by activated pancreatic stellate cells in the tumor microenvironment (TME)(*14, 15*). Elevated collagen is a hallmark of several other desmoplastic solid tumors, including breast carcinoma where it is associated with higher stiffness, hyperproliferation, increased invasion and metastasis (*16–19*). In the context of invasion, not only the amount of collagen, but also the architecture of the fibers in the TME are vitally important. In particular, several distinct collagen organization patterns, termed tumor-associated collagen signatures (TACS), have been identified in breast tumors with important implications in disease progression (*20, 21*). Among them were TACS-2 and TACS-3, both comprising of organized collagen fibers in the periductal space. For TACS-2, collagen fibers of variable density are mainly organized approximately parallel to either the ductal or carcinoma in situ boundary, around carcinoma cell clusters within the tumor mass, or at the tumor boundary. For TACS-3 the fibers orient perpendicularly to the cell clusters or ductal boundary, or result in aligned collagen regions throughout the tumor mass in later stages, often providing a conduit for carcinoma cell invasion (*20, 22*). Consequently, due to contact guidance where cells utilize anisotropy from aligned ECM fibers to orient and migrate along single fibers, TACS-3-like aligned collagen patterns lead to increased focal and local invasion and metastasis (*20, 22*) and correlate with worse survival in human patients (*21*). Yet, while recent studies suggest the presence of TACS architectures in PDA (*10, 23*), the prevalence of TACS in PDA, particularly relative to disease stage and early dissemination remains largely unexplored. Thus, we hypothesized that TACS or TACS-like collagen architectures are involved in early dissemination and invasion in PDA.

Here, we employ an autochthonous mouse model of PDA, highly faithful to the human disease (*24*) and expressing carcinoma cell-specific fluorophores (*10*), as well as human PDA samples to define a key link between early dissemination in PDA and periductal collagen organization. We utilize integrated multiphoton excitation (MPE) and second harmonic generation (SHG) imaging on archival and live tumor samples to characterize the prevalence of TACS in PDA and demonstrate that periductal collagen patterns drive carcinoma cell dissemination. We further identify these patterns in pancreatitis and inflamed “normal” adjacent tissue, suggesting that precancerous fibroinflammatory disease can precondition and prime the stroma for early dissemination prior to transformation. Moreover, we identify focal adhesion kinase (FAK), a key mechanotransduction kinase molecule, as a regulator of the processes of ECM-guided extrusion from ductal epithelium and subsequent invasion, and observe that blocking FAK function reduces the efficiency of single cell dissemination and metastasis in PDA.

## Results

### Evolution of collagen organization in PDA

To characterize the deposition and architecture of fibrillar collagen with disease progression in PDA, we utilized SHG imaging for label-free detection of collagen from live and archival murine and human tissue samples (Fig. 1). Multiple large (~1mm^2^), human patient biopsy sections on a tissue microarray (complete with tumor straging and grading information) were imaged at high resolution by multiphoton microscopy to generate simultaneous SHG and MPE of endogenous cellular fluorescence (Supplementary Fig. 1a), enabling quantification of fibrous collagen from the SHG signal (Supplementary Fig. 1b), while also mapping the localization of the collagen fibers with respect to tissue architecture. In normal pancreata, little collagen was observed, largely concentrated around the few ducts interspersed in tissue otherwise dominated by acinar cells (Fig. 1a, e; note no to low collagen surrounds acinar cells (see magnified region #) in contrast to robust collagen surrounding ductal structures (see magnified region ##)). Notably, cancer-adjacent normal (CAN) regions showed elevated levels of collagen around ductal structures and in some regions between acinar clusters (Fig. 1b, e; note acinar regions can possess both collagen (main image) or no collagen regions (see magnified region #)), suggesting regional activation of a fibroinflammatory stromal response in CAN tissue, consistent with the intimate link between fibroblast activity and collagen deposition in PDA development (*3, 25*). This may contribute to epithelial dysfunction in adjacent regions (e.g. acinar dropout, ductal hyperplasia, or acinar to ductal metaplasia (ADM)) and/or prime adjacent regions for invasion. Moreover, consistent with the desmoplastic response associated with PDA, as expected, robust fibrous collagen was ubiquitous in the periductal areas around pancreatic intraepithelial neoplasia (PanIN) lesions, which are histologically well-defined precursor to invasive ductal pancreatic adenocarcinoma (Fig. 1c, e). Elevated collagen remained pervasive throughout disease progression to mature PDA (Fig. 1d, e).

**Figure 1:**
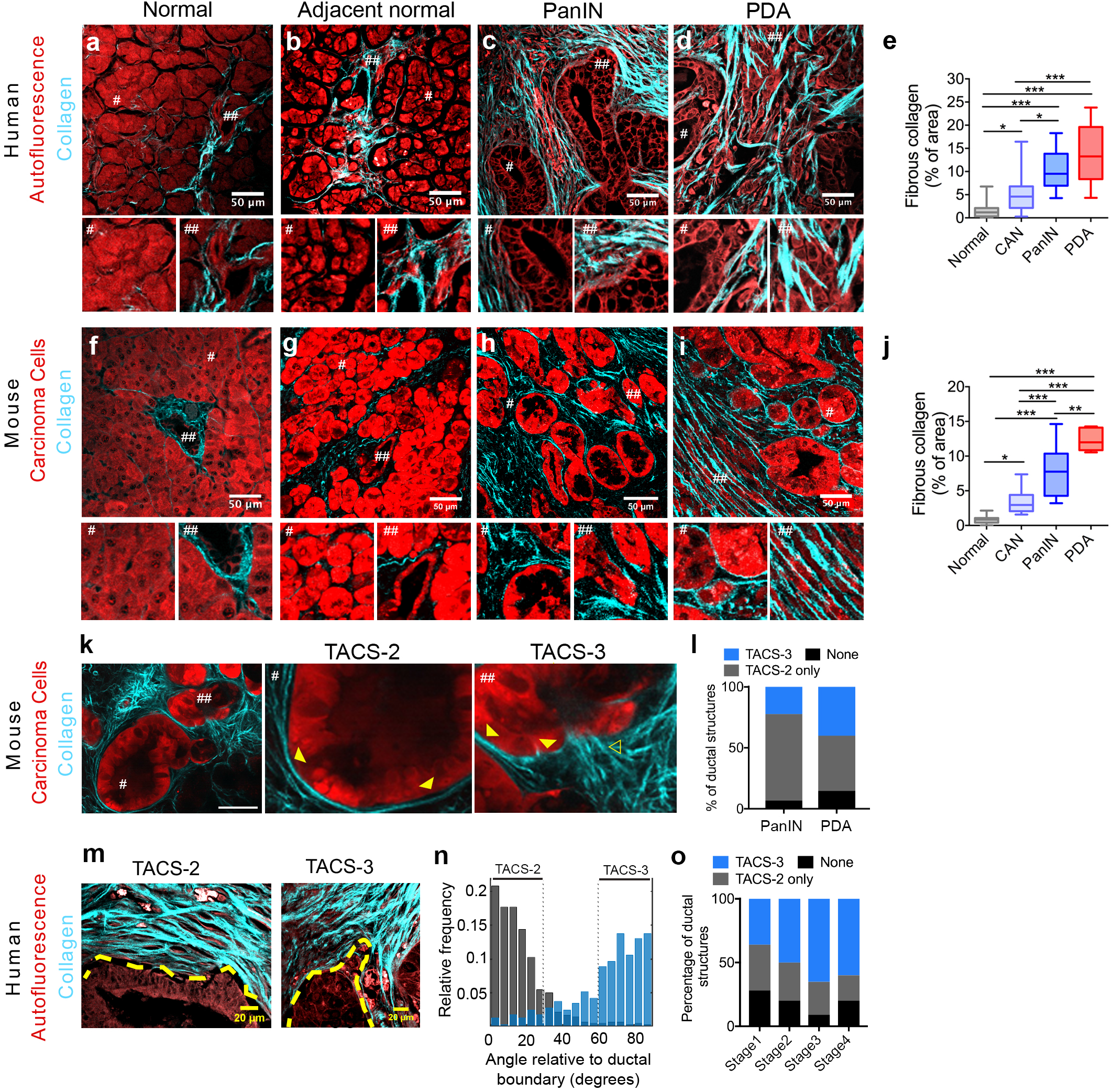
Fibrous collagen architectures in PDA: (**a-d**) Combined multiphoton excitation (MPE) and second harmonic generation (SHG) of human biopsy samples (from a human PDA tissue microarray; TMA) showing tissue architecture by endogenous fluorescence (autofluorescence) and fibrous collagen by SHG in (a) normal, (b) cancer-adjacent normal, (c) PanIN and (d) mature PDA (# and ## are 2X magnifications of the indicated regions). (**e**) Collagen quantification for the data shown in a-d (n=10-21 fields of view across n≥3 patient biopsies). (**f-g**) Combined MPE/SHG imaging and analysis using *KPCT* or *KPCG* mouse models of PDA for (f) normal, (g) cancer-adjacent normal, (h) PanIN and (i) well-differentiated PDA. Note, all tdTomato or zsGreen fluorescence is pseudo-colored red for consistency of identifying carcinoma cells. (**j**) Collagen quantification for the data shown in f-i (n=5-12 fields of views across n≥3 mice). (**k**) Example of collagen architectures in early stage disease (1.5mo) in a *KPCT* mouse (# and ## are 3X magnifications of the indicated regions). Outlined and solid arrowhead point to TACS-3 and TACS-2, respectively (Note that TACS-3 positive ducts are also TACS-2 positive in other regions of the duct). (**l**) Frequency of quantified TACS (per duct) associated with PanIN lesions and well-differentiated PDA genetically engineered mice. Note, the vast majority of ducts are TACS2 positive, and TACS-3 positive ducts are also TACS-2 positive in different regions (see Supplementary Figure 1c). (**m-n**) Representative TACS-2 and TACS-3 positive ducts in well-differentiated human PDA (i.e. around easily identifiable ductal structures) showing parallel and perpendicular alignment of fibrous collagen, with yellow dashed lines indicating ductal boundaries (m), quantified by CT-FIRE (n). (**o**) Frequency of TACS-2 and TACS-3 associated with ductal structures in well-differentiated regions from different stages of human disease (from pathology staged and graded TMA) showing increased prevalence TACS-3 with more advanced disease. Scale bars = 50 μm (a-d, f-I, and k) and 20 μm (m), box-whisker plots show min to max with median and interquartile range for e and j; *p<0.05, **p<0.01, ***p<0.001 by the non-parametric Kruskal-Wallis test and Dunn’s multiple comparison test.

To complement analyses from human patient biopsies, we employed the genetically engineered *Kras^LSLG12D/+^;p53^LSL-R172H/+^;Pdx-1-Cre* (*KPC*) mouse model that faithfully replicates the human disease, including the stromal response associated with PDA (*5, 24*) and provides an excellent tool to examine collagen organization during PDA progression. Here, we utilized fluorescent reporter *KPC* mice, namely *KPC;ROSA26^LSL-tdTomato/+^* (*KPCT*) or *KPC;ROSA26^LSL-ZsGreen1/+^*(*KPCG*; pseudo-colored red throughout the work here for consistency unless otherwise noted) and performed similar analyses of the localization and distribution of collagen (Fig. 1f-j). Consistent with the human data, normal murine tissues possess minimal stromal collagen while CAN regions show regions of elevated collagen deposition around ducts that extend into the periacinar space (Fig. 1g, j). Likewise, histologically “pre-invasive” PanIN regions were highly collagenous with robust periductal collagen observed around the majority (>95%) of the ductal structures (Fig. 1h, j). Overall, these data from human and murine PDA samples demonstrate biased collagen localization around ductal structures in normal pancreata, elevated collagen levels in adjacent normal regions, and more robust collagen in early PanIN lesions and PDA.

Since elevated collagen levels in breast carcinomas are organized into specific architectures like TACS-3, which guide local invasion ((*16, 20, 22*); Supplementary Figure 1c), and both TACS-3 architectures (*5, 10, 23*) and early invasion (*11*) have been observed in PDA, we hypothesized that such architectures may also be prevalent in very early (histologically “pre-invasive”) pancreatic cancer and play a key role in early invasion. This prompted us to characterize the periductal collagen organization in histologically early disease from *KPC* mice. Analysis of collagen architecture in the periductal space demonstrates that ducts are TACS-2 positive (Figure 1f-i). That is, analysis of SHG images shows parallel collagen distributed around ~0° (defined as +/− 30° similar to our previous reports (*20, 26*)) relative to the epithelial boundary (Supplementary Fig. 1c, see also magnified regions ## of ducts in Figures 1f and g, and TACS-2 organization shown in magnified regions # of PanINs and well-differentiated PDA regions in Figures 1h and i). We note that in pancreatic disease, collagen is often less tightly bound around ductal structures than we previously observed when defining TACS-2 architectures observed in mammary carcinomas (*20*). Furthermore, analysis of TACS-2-positive ducts from early disease in ~1.5 month old mice (preceding frank tumor formation) indicates that many of the ducts are also TACS-3 positive (defined as collagen aligned perpendicular, ~90 +/− 30° similar to our previous reports (*20, 26*), relative to the epithelial boundary; Supplementary Figure 1c-e). That is, regions of the duct are positive for TACS-2 while adjacent regions of the ducts are positive for TACS-3 (Figure 1k, see also TACS-2 and TACS-3 architectures surrounding PanIN lesions in Figure 1h and Supplementary Figures 1e), demonstrating that collagen surrounding “pre-invasive” PanIN lesions possess architectures that are known to promote invasion of pancreatic carcinoma cells (*10*). We note that both TACS-2 and TACS-3 architectures also exist in well-differentiated PDA regions (i.e. right half of Figure 1i), demonstrating that such aligned collagen architectures are maintained near ductal structures associated with more advanced disease. While a significant fraction of the ductal structures in early disease present with TACS-3 regions, they are slightly more frequent in advanced well-differentiated PDA as compared to PanIN lesions (Fig. 1l).

In agreement with murine PDA data, both TACS-2 and -3 are found surrounding ductal structures in human pancreatic cancer. TACS-2 is again robustly observed around the vast majority of ductal structures in early and well-differentiated PDA regions (Fig. c, d). Frequently, TACS-3 is observed locally, often associated with invaginations and regions of irregular ductal boundaries (Fig. 1m-o). Indeed, similar to findings in *KPC* mice, quantitative analysis of fiber orientations using a Curvelet Transform of the SHG signal demonstrates TACS-2 (again defined as parallel collagen distributed around ~0° +/− +/− 30° relative to the ductal boundary) or TACS-3 (again defined as perpendicular collagen distributed around ~90° +/− 30° to the ductal boundary) periductal collagen organization (Fig. 1n). Notably, in human samples with Stage-I to Stage-IV disease, the presence of TACS architectures was not limited to a particular disease stage (Fig. 1o). However, consistent with invasive breast carcinoma (*20, 21*), there tends to be a higher prevalence of TACS-3 positive ducts in more advanced disease (Fig. 1o). Yet, a substantial proportion (~40%) of ductal structures in Stage-I patients also present with TACS-3 (Fig. 1m), demonstrating that these conduits for invasion are present at early stages and remain prevalent in more advanced PDA. Thus, our findings that robust TACS-3 surrounding ductal structures emerges with histologically “pre-invasive” disease and is maintained through advanced well-differentiated disease, combined with our understanding of the established role of aligned collagen in promoting highly directed carcinoma cell invasion (*10, 22, 27–31*), motivated us to further evaluate the influence of collagen alignment on early dissemination of pancreatic carcinoma cells.

### Single cell dissemination along local periductal collagen architectures

Rhim and colleagues (*11*) demonstrated that invasive dissemination can begin early in PDA development, even prior to detection of histologically malignant disease and frank tumor formation, resulting in single cells in the tumor stroma and DCCs in the blood of 8-week old *KPC* mice (*11*). This is strikingly consistent with the timeline of robust collagen deposition and establishment of TACS architectures observed here and therefore led us to investigate whether such collagen patterns are key enablers of early dissemination. Furthermore, we note that extrusion of cells from the epithelium and into the lumen (i.e. apical extrusion) is a normal process for epithelium turnover, where apically extruded cells undergo apoptosis via anoikis (*32, 33*). However, basal extrusion has also been described in normal (*32*) as well as malignant (*33, 34*) ductal development and maintenance, and would precede invasion through the stroma for early dissemination from histologically preinvasive lesions. Therefore, to characterize carcinoma cell extrusion from ductal structures, we performed combined MPE and SHG imaging over multiple regions in several pancreatic tumor sections from a number of *KPCT* and *KPCG* mice. Using ZsGreen1 or tdTomato to identify carcinoma cells (from *KPCG* and *KPCT* mice, respectively) colocalized with a nuclear stain and SHG for collagen, we identified different stages of extrusion associated with distinct collagen patterns. Partially disseminated cells were found extensively around ductal structures in PanIN lesions and well-differentiated PDA and were, perhaps surprisingly, associated with both TACS-2-like structures and TACS-3 (Fig. 2a, b). Strikingly, these partially disseminated cells at the ductal boundary, while still attached to the main duct, were aligned in the direction of local collagen organization (Supplementary Figure 2a,c), i.e. demonstrating contact guidance (*10, 28*). For PanIN lesions, ~53% of the partially extruded cells on average were parallel and about 28% perpendicular to the ductal boundary (Fig. 2c); that is, associated with pancreatic TACS-2 or TACS-3 periductal collagen fiber architectures, respectively. This demonstrates that pancreatic carcinoma cells can enter the less constrained (i.e. looser than the tighter TACS-2 architectures observed in mammary carcinomas (*20*)) TACS-2 architectures observed in PDA, and likely undergo directed motility along the collagen around the ductal structure. In contrast to early disease, partially extruded cells were equally likely to be on TACS-2 or TACS-3 architectures in advanced well-differentiated disease (Fig. 2d). This difference is likely the result of the increased frequency of TACS-3^+^ ductal structures in more advanced disease as compared to early PanIN lesions (Fig. 1l, o). Importantly, similar partial dissemination associated with periductal collagen was observed in well-differentiated ductal regions of human PDA sections, where a pan-Cytokeratin stain was utilized to visualize carcinoma cells (Fig. 2e). Such partially delaminated cells likely represent the early stages of extrusion and their alignment with the surrounding collagen is indicative of the close association between the ductal epithelium and periductal collagen during this process.

**Figure 2:**
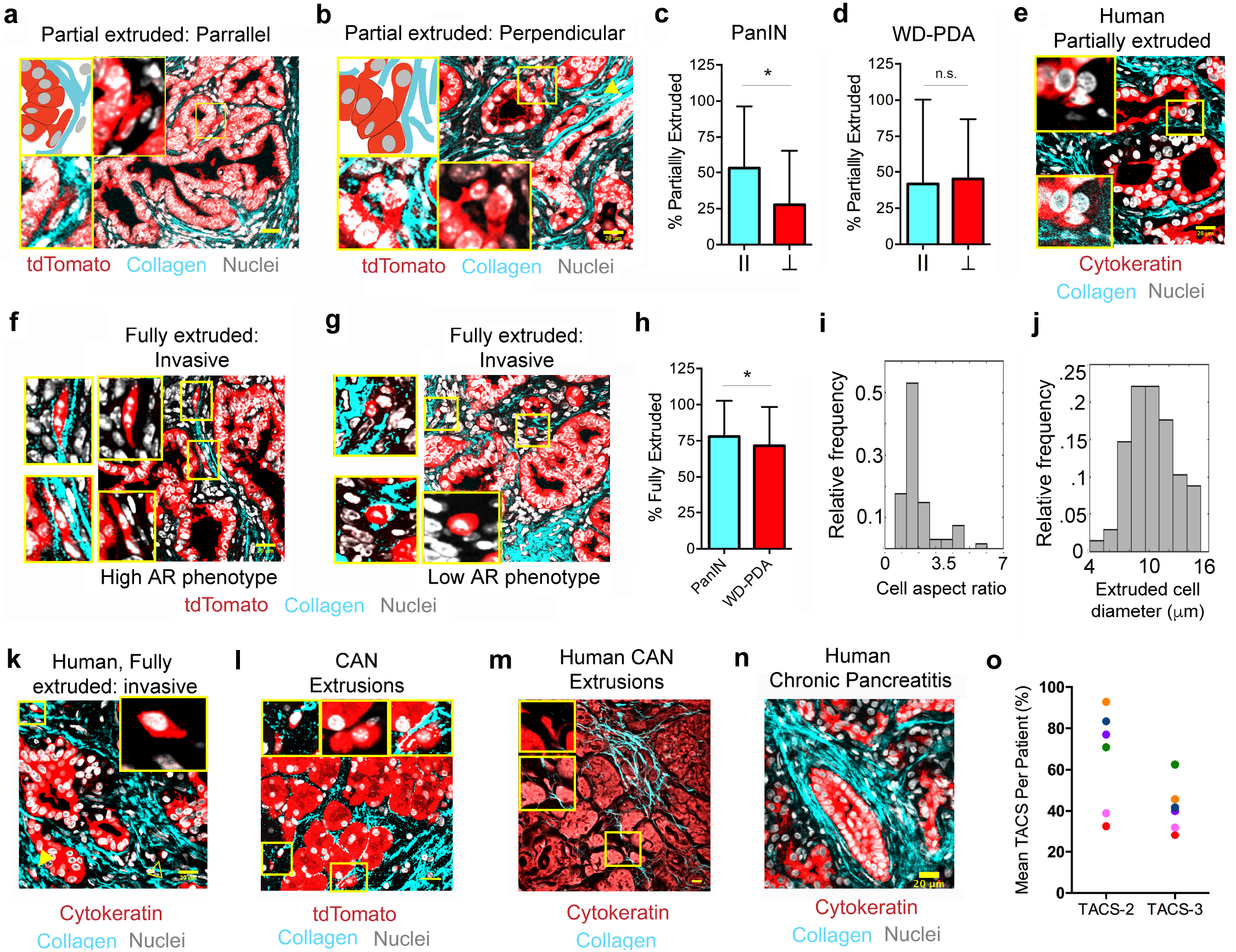
Characterization of PDA cell extrusion into periductal collagen architectures: (**a,b**) Schematic (*top left* boxes) and micrographs of partial extrusion of carcinoma cells as observed in *KPCT* or *KPCG* tumor sections showing cells extruding into (**a**) TACS-2 and (**b**) TACS-3 architectures in the periductal stroma. *Yellow boxes* represent zoomed in images showing either all three channels (*bottom left*) or only the fluorescent reporter and nuclei channels (*insets*). *Yellow arrowhead* (in b) points to an additional cell that is fully extruded and aligned to the collagen fibers. (**c,d**) Quantification of frequency and types of partial cell extrusion in (**c**) PanIN and (**d**) well-differentiated PDA (WD-PDA); II and ⊥ indicates parallel TACS2 and perpendicular TACS3 architectures, respectively; data are mean +/− SD. (**e**) Partial extrusion observed in human well-differentiated PDA samples. *Yellow boxes* represent the zoomed region showing either all three channels (*bottom inset*) or only the fluorescent reporter and nuclei channels (*top inset*). (**f-h**) Fully extruded carcinoma cells invading through the stroma (f,g) were observed at similar frequency in PanIN and well-differentiated PDA (h) and cells along collagen fibers displayed either an (f) elongated and oriented phenotypes (high AR phenotype) or (g) a more rounded phenotype with smaller cell diameters (Low AR phenotype); *Insets* represent zoomed in images, identified with yellow boxes, showing only the fluorescent reporter and nuclei channels. (**i.j**) Morphometric analysis of extruded cells shows a large range of (i) aspect ratios and (j) diameters (high and low aspect ratios cells are represented in (f) and (g) respectively. (**k**) Full extrusion of invading cells from ductal structures was observed in human well-differentiated PDA samples; *Insets* represent zoomed in images identified with yellow boxes showing the fluorescent reporter and nuclei channels, *solid yellow arrowhead* (in k) points to an additional, aligned, fully extruded cell and the *outlined yellow arrowhead* points to one that is rounded and non-aligned. (**l,m**) Cancer adjacent “normal” regions from *KPC* mice or human disease showing basal extrusions associated with collagen fibers. Insets show magnified images of cells marked with yellow boxes, either with all three channels or with the fluorescent reporter and nuclei. (**n**) Collagen architecture surrounding ducts in human chronic pancreatitis (CP). (**o**) Frequency of TACS-2 and TACS-3 in association with ductal structures in human CP (Analysis is from 8-33 fields of view from two biopsies per patient from 6 patients with human CP; note, every patient sample presented with ducts that were TACS-3-positive). Scale bars = 20 μm, n >50 fields of view across 11 mice (c), n >20 fields of view across 6 mice (d and h), n=68 cells across 12 mice for (i) and (j); data are mean +/− SD.

In addition to partial extrusion, we observed an abundance of fully extruded, invasive, cells in the periductal space of PanIN lesions and well-differentiated PDA (Fig. 2f-h, Supplementary Figure 2a,b), consistent with previous reports identifying carcinoma cells in the stroma during early disease (*11, 35*), and with carcinoma cells principally following collagen aligned architectures due to contact guidance (Figure 2, Supplementary Figure 2a). To confirm these findings we imaged live tumors from *KPCT* mice with combined MPE and SHG microscopy, which also allows for z-stack imaging around dispersed cells to confirm the presence of extruded invasive cells (Supplementary Figure 2c). Consistent with findings in fixed samples, extruded cells are present and the majority aligned along TACS-2 and TACS-3 architectures, demonstrating contact guidance. Thus, as we observe TACS-2 architecture connecting to TACS-3 regions (Supplementary Figure 2d) we speculate that carcinoma cells extruded into TACS-2 regions migrate along collagen around ducts until encountering a TACS-3 exit point or do not easily leave the periductual space since migration perpendicular to dense collagen is profoundly less efficient and in many cases extremely rare (*10, 22, 27, 36, 37*). Furthermore, quantitative analysis of the phenotype of single cells in the periductal space demonstrates that invading cells assume two distinct morphologies. While the majority (~70%) of the cells are elongated and aligned to the orientation of the periductal collagen (Fig. 2f), a distinct proportion of more rounded, single cells also exist along collagen in the stroma (Fig. 2g), consistent with our previous findings that both phenotypes undergo directed migration from contact guidance cues (*10*). This is further supported by morphological analysis revealing a large range of aspect ratios (AR) and small cell diameters for fully extruded cells (Figs. 2i, j). A significant proportion of cells (~30%) indeed have an aspect ratio of less than 1.5 and likely represent a distinct phenotypic population in addition to the elongated, more phenotypically mesenchymal-like single cells, which is consistent with previous observations of heterogeneous breast carcinoma cell phenotypes during 3D migration from contact guidance (*27*) and distinct phenotypes and phenotypic plasticity reported in PDA (*11, 38, 39*). Moreover, these two separate phenotypes can be observed in fully extruded cells in human patient samples, often existing in the same local region (Fig. 2k), further supporting that analysis of cell extrusion in the *KPC* model is highly relevant to human disease, and suggesting that phenotypically distinct contact-guided subpopulations are capable of invasion leading to metastasis in PDA.

In contrast to normal tissues that have very little to no collagen around acinar cells and where we did not observe any basal extrusions in either murine or human pancreata (Supplementary Figure 2b), and similar to PanIN lesions and well-differentiated PDA, extrusion events were present in CAN regions adjacent to disease in murine and human tissue, albeit at a lower frequency than observed with PanINs (Figs. 2l,m, Supplementary Figure 2b). It is striking that even in CAN regions, which have lower collagen content than PanIN lesions, disseminated cells were almost always associated with regions of peri-acinar or periductal fibrous collagen (Figs. 2l,m). This suggests that the fibro-inflammatory response occurring in tissue directly adjacent to disease may aid the extrusion process, priming the stroma in adjacent regions to facilitate disease spread prior to infiltration of frank malignant disease into that these regions. Therefore, we explored TACS architectures in pancreatitis, which represents a strong risk factor for development of PDA and can be a precursor to malignancy (*40*). Importantly, pancreatitis can display a PDA-like desmoplastic response with an accumulation of straight and thick collagen fibers (*23*). Indeed, analysis of biopsies from 6 human patients with chronic pancreatitis revealed an abundance of TACS-2 and TACS-3 associated with ductal structures (analysis from 8-33 fields of view from two biopsies per patient; Fig. 2n, o). Consistent with our findings in pancreas cancer, approximately 32-93% of ductal structures were TACS-2 positive and ~28-62% were positive for both TACS-2 and TACS-3 (similar to pancreatic cancer, ducts that are TACS-3+ are also TACS-2+ in different regions of the duct; Fig. 2o). Remarkably, every pancreatitis patient sample presented with ducts that were TACS-3 positive. However, extrusion events were only infrequently observed relative to findings in PanIN lesions and well-differentiated PDA. This suggests that transformed carcinoma cells are more robustly primed to basally extrude when presented with aligned ECM architectures, as compared to untransformed epithelial cells, or that the desmoplastic phenotype is not as robust as malignant regions, or both, raising important questions regarding the cooperative cell-intrinsic and extrinsic mechanisms governing dissemination. Moreover, these data indicate that fibroinflammatory diseases that may precede PDA, such as pancreatitis, can result in stromal ECM architectures primed to facilitate disease spread from the earliest onset of disease. This suggests that therapeutic strategies to resolve fibroinflammation, such as stroma-targeting antifibrotic therapies (*4, 41*), may be beneficial in disrupting ECM architectures conducive to disease spread, thus impeding development of early DCCs in patients who later develop PDA.

### Dynamics of single cell extrusion and contact guided migration in stromal collagen

Given the role periductal collagen plays in facilitating basal cellular extrusion, we speculated that TACS located distal to ductal structures also influences cancer cell dissemination through the stroma. Indeed, fully extruded cells in both murine and human samples were found to be colocalized with aligned collagen (Fig. 2f-h, 2k, Supplementary Figure 2a,c). This suggests that TACS not only aid in the dispersal of these cells from the epithelium during basal extrusions, but also guide the extruded cells as they invade through the stroma. Indeed, we have previously demonstrated that carcinoma cells, including those of the pancreas, are strongly directed to migrate along aligned ECM via contact guidance with minimal motility lateral to ECM fiber alignment (*10, 22, 27–29*). Live MPE/SHG imaging of pancreatic tumor slices again confirmed the prevalence of extruded cells in the periductal space around both PanIN lesions and in well-differentiated PDA (Fig. 3a, Supplementary Figure 2). Moreover, to further confirm that the features observed were not an artifact of tissue slicing, i.e. the extruded cells were indeed discrete entities and not part of a larger ductal network connected in other planes, we reconstructed the 3D volume around entire ductal structures using multiphoton imaging of live early disease tumor samples, and again confirmed the existence of single carcinoma cells in the stroma (Fig. 3b, Movie 1). Interestingly, long-term imaging of ductal structures also revealed dynamic rearrangement and rotational movement of the epithelium, a phenomenon known as coherent angular motion (Movie 2), which has been previously observed *in vitro* in mammary acini (*42, 43*), and is thought to be sustained, in part, by cell division (*43*). Our observation of this phenomenon in live pre-malignant and malignant tissue further demonstrates the dynamic nature of epithelial cells in organized ductal structures. These cellular dynamics are perhaps intricately tied to the process of cell extrusion as a mechanism to maintain epithelial tissue homeostasis in the face of aberrant cell division, as in cancers (*33*).

**Figure 3:**
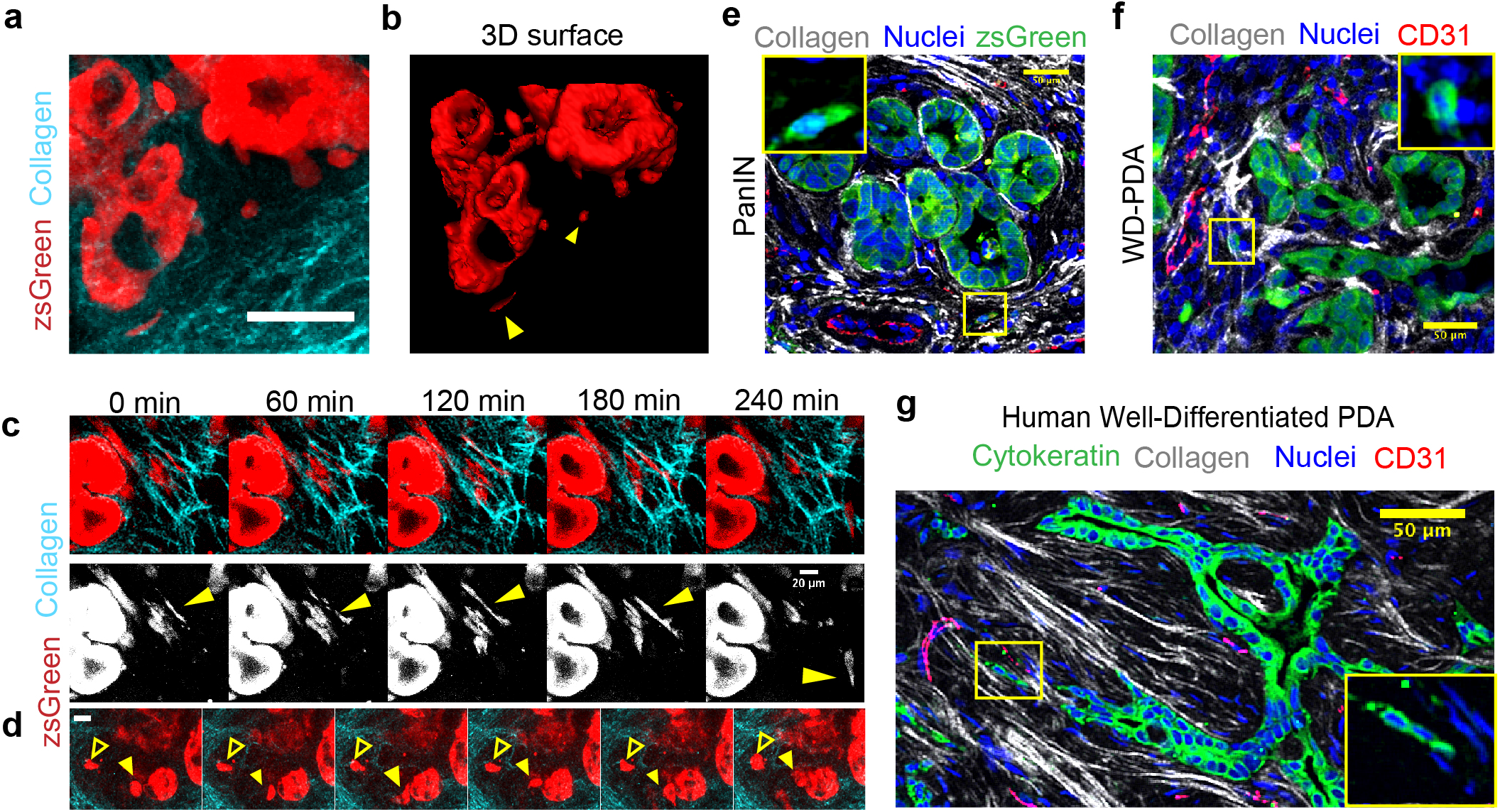
Live imaging reveals dynamics of cell extrusion and invasion in early stage disease: (**a,b**) Live 3D imaging of PanIN lesions in a *KPCG* mouse (zsGreen pseudo-colored red) showing extruded cells invading through organized collagen in the periductal space by a (a) maximum intensity projection of ~100μm z depth and (b) 3D surface rendering of the same region demonstrating that the extrusion events observed in tissue slices were indeed single cells, not attached to the main duct at other z planes; *yellow arrowheads* point to extruded cells - see also Movie 1. (**c**) Time-lapse montage from tracking single, aligned cells in periductal collagen by live MPE/SHG imaging of a *KPCG* tumor, *yellow arrowhead* indicates cell of interest that migrates rapidly along a collagen fiber - see also Movie 4. (**d**) Time-lapse montage showing the dynamics of a partially and fully extruded cell in the same field of view in a *KPCG* tumor, *yellow solid arrowhead* points to a partially extruded cell, still connected to the underlying ductal structure and *yellow outline arrowhead* points to a fully extruded cell in the stroma - see also Movie 5. (**e, f**) Immunofluorescence micrograph of *KPCT* tumor sections stained with RFP (shown in green), CD31 (red) and DRAQ5 (blue), demonstrating single extruded cells interacting with aligned periductal collagen (white) directed to blood vessels in (e) PanIN lesions and (f) well-differentiated PDA; *inset* shows magnified region enclosed by the yellow box displaying only the fluorescent reporter and nuclei channels. (**g**) Fluorescence micrograph of human PDA stained with CD31 (red), DRAQ5 (blue), and Cytokeratin (green) showing aligned extruded cells following collagen tracks (white) leading to a blood vessel; *inset* shows magnified region enclosed by the region enclosed by the yellow box displaying only the fluorescent reporter and nuclei channels. Scale bars = 20 μm (c, d) and 50 μm (a, b, e-g).

Furthermore, consistent with static imaging, analysis of live cell dynamics revealed motile cells that have invaded into the stroma. We observed a minor population that are directed by collagen architecture and showed transient switching between more rounded and more elongated phenotypes (Movie 3), consistent with cell aspect ratio analysis (Figure 2). Concurrently we also observe the larger population of highly elongated cells (i.e. high AR) associated with aligned periductal collagen, rapidly migrating along the collagen tracks (Fig. 3c, Movie 4). We note that these two populations are reminiscent of the distinct phenotypes and plasticity that have been separately associated with invasion and metastasis in PDA (*11, 38, 39*) as well as the contact guidance response of breast carcinoma cells where we observed a spectrum of phenotypic heterogeneity, from epithelial to those that have undergone epithelial-to-mesenchymal transition (EMT), and robust phenotypic switching (*10, 27*). Furthermore, through live imaging we observe cell extrusion from a ductal structure alongside already extruded cancer cells in the periductal stroma (Fig. 3d, Movies 4 and 5), providing additional compelling evidence demonstrating basal extrusions from ductal structures in PDA. Thus, the combined data suggest the potential for a consistent stream of pre-metastatic dissemination from ductal epithelium, which may not be solely dependent on a stable EMT phenotype but inevitably contribute to the early metastatic spreading associated with pancreatic cancer.

In line with our conclusions of that early extrusion events leading to invasive cells in the stroma contribute to early metastatic spread we note that reports indicate that PDA carcinoma cells can be present in circulation during histologically pre-invasive disease (*11*) and that in other desmoplastic systems such as in breast carcinoma, pioneer metastatic cells utilize bundles of aligned collagen as highways to escape towards blood vessels (*44*), often further guided by macrophages (*45*) in the tumor stroma. Therefore, we visualized the localization of TACS and extruded cells with respect to CD31^+^ blood vessels in archival tissue samples. Carcinoma cells in the stroma were frequently observed in close association with blood vessels in the stroma (Fig. 3e-g; Supplementary Figure 3). In *KPC* mice, extruded single cells were present on aligned collagen fibers leading to open blood vessels (which are less frequent relative to the larger fraction of collapsed vessels in PDA, (*5, 6, 41*)) in both PanIN lesions as well as more mature well-differentiated disease (Fig. 3e,f; Supplementary Figure 3). Importantly, we confirmed these findings in human tissue samples using pan-cytokeratin to mark cancer cells (Fig. 3g). Again, aligned collagen fiber tracks, punctuated with single or streams of multiple carcinoma cells, were observed leading to CD31^+^ blood vessels (Fig. 3g; Supplementary Figure 3). It is important to note that while a large percentage (~75%) of blood vessels in mature PDA are collapsed and non-perfused (*5, 6*), those in the early stages of the disease have a much higher likelihood of being open (*41*) and thereby likely provide a more clear passage for these constantly extruding cells to escape and enter the bloodstream. Taken together, these data strongly suggest that the phenomenon of early extrusion and invasion along organized collagen fibers plays a direct role in early and extensive metastasis observed in PDA. We also note that this program is prevalent through all stages of disease, actively and continually promoting disease spread, suggesting that disrupting this process may be a viable strategy to slow or halt disease progression during therapeutic interventions to combat already established primary and metastatic disease. However, while the intra and intercellular dynamics during basal cell extrusion from ducts has been explored (*33, 34, 46*), how the ECM affects this process is unexplored, prompting us to delve into its biophysical and molecular mechanisms.

### FAK-dependent mechanotransduction enables collagen-guided cell extrusion and invasion *in vitro*

The data presented thus far point to periductal collagen architectures as well as cell-intrinsic properties as key factors guiding single cell dissemination from ductal structures in PDA. Extending our understanding of single cell contact guidance (*10*) to multicellular assemblies in ductal structures transitioning to collective or single cell invasion exposes an interesting duality between cell-cell dynamics and cell-matrix interactions from stromal collagen. A cell at the interface is presented with multiple, counteracting cues to either remain in its cohesive state in the epithelium or break away from its native structure and follow ECM alignment. Indeed, we note that primary PDA cells derived from *KPCG* or *KPCT* mice can display strikingly contrasting morphologies in 3D collagen matrices that have fiber alignments versus 3D Matrigel, the latter representing a more basement membrane-like environment (Supplementary Fig. 4a). While cells tended to cluster together in tight epithelial-like structures in Matrigel, they form more tubular, spindle-shaped, protrusive structures in 3D when embedded in a fibrous collagen matrix (Supplementary Fig. 4a). This striking difference suggested to us that cells at the interface of a basement membrane-bound epithelial structure and surrounding fibrous collagen matrix experience uniquely counteracting cues that may lead to the extrusion phenotype observed in ductal epithelia in presence of organized periductal collagen. To dissect this process, we harnessed microfluidic technology to generate primary PDA cell-derived epithelial organoids in a high throughput system. These organoids could be subsequently embedded into an ECM whereby epithelial cells encounter similar counteracting cues of cell-cell interactions with basement membrane and aligned collagen (Fig. 4a-f; Supplementary Figure 4b,c). Briefly, droplets of Matrigel were generated by an oil-aqueous interface in a microfluidic flow focusing device (Fig. 4a) and layered with primary *KPCT or KPCG* cells (Fig. 4b). The cells remain organized in a ring around the Matrigel drop, rarely invading into the droplet and resembling a ductal cross-section, as evidenced by 3D imaging (Fig. 4b-e). These microtissues were cultured in individual agarose microwells to allow them to evolve into robust epithelial organoids over several days where they still retain basement membrane (Fig. 4d,e) and subsequently embedded in matrices where the interaction of individual organoids with the surrounding ECM could be studied (Fig. 4f; Supplementary Figure 4b,c). Indeed, within 24h of embedding, cancer cells began to extrude and disseminate along collagen fibers (Fig. 4g), where collective invasions and single cell extrusions to invasion were observed (Movies 6-8). In contrast, organoids embedded in Matrigel largely retain a rounded ductal morphology (Supplementary Figure 4b,c). Importantly, in collagen matrices the features of extruding regions are highly reminiscent of ductal outgrowths and collective and single cell dissemination associated with TACS-3 observed in murine and human PDA samples (Figures 1 and 2, Supplementary Figures. 2c, 4d). Further, rapid extrusion led to extremely robust invasion under control conditions over 3 days (Figure Supplementary Fig. 4e), consistent with findings with mammary epithelial acini (*47*). These data indicate that our novel *in vitro* system, particularly at early time points, can recapitulate key features of the extrusion to invasion process and provides a high throughput controllable platform in which to investigate the molecular mechanisms guiding epithelial extrusion in PDA.

**Figure 4:**
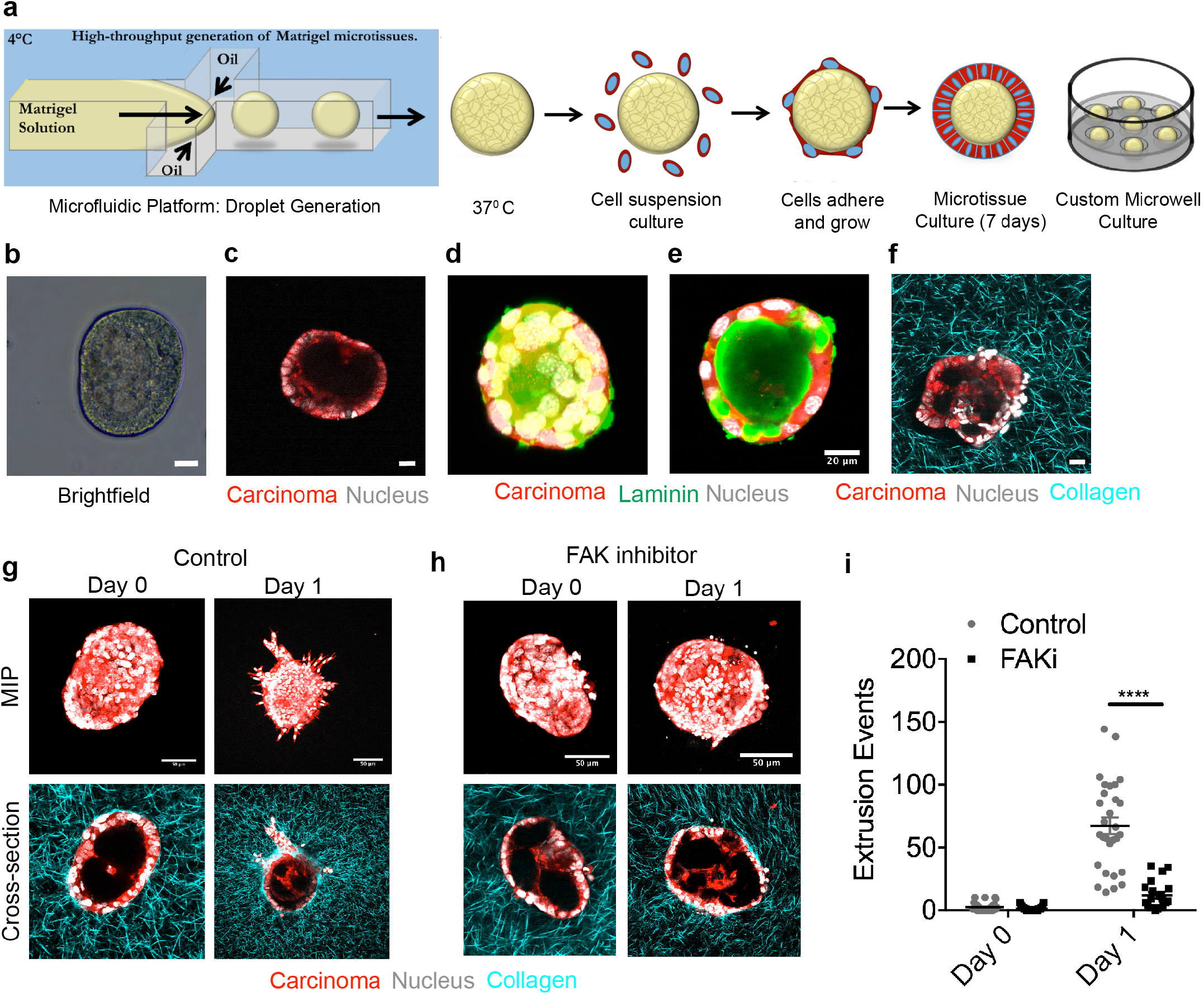
FAK inhibition abrogates collagen fiber guided PDA cell extrusion *in vitro*: (**a**) Schematic demonstrating our *in vitro* platform to mimic tumor cell extrusion from ductal structures into surrounding collagen. Matrigel droplets generated in a microfluidic device by orthogonal flow of Matrigel in an oil based medium, before coating them with primary *KPCG* or *KPCT* cells. Coated droplets are cultured individually in custom-fabricated microwells for several days before embedding in a collagen matrix. (**b,c**) Brightfield (b) and multiphoton excitation microscopy image (c) of a formed microtissue. (**d,e**) Maximum intensity projection (MIP, d) and cross-sectional (e) MPE immunofluorescence micrographs show a typical droplet structure resembling the cross-section of a duct with basement membrane. (**f**) MPE/SHG imaging of a droplet embedded in collagen matrix. (**g-i**) Control (g) and FAK inhibitor (h) treated conditions, showing significant invasion in the control group at Day 1, which is completely abrogated by inhibition of FAK, as quantified by single cell extrusion analysis in (i); n=20-30 droplets per condition from n=2 experiments, Data are mean +/− SEM; ****p<0.0001 by ordinary 2-way ANOVA and Sidak’s multiple comparison test. Scale bars=20μm (b-f) and 50μm (g, h).

The fact that embedding these epithelial structures within collagen matrices is sufficient to induce extrusion and invasion is consistent with the conclusion that this dynamic process is guided by structural cues in the periductal ECM. In response to this extrinsic factor, we investigated cell-intrinsic signaling that may modulate cell extrusion. Recent work has shown that in stiff environments contact guidance is orchestrated by anisotropic forces arising from constrained focal adhesion maturation on discrete collagen fibers and that intercellular forces can counteract such directional cell-ECM forces (*10, 48*). A corollary to this theory suggests that a carcinoma cell interacting with organized periductal collagen may experience sufficient anisotropic forces to overcome strong intercellular forces and extrude out of the epithelium. Thus, the intercellular force generation machinery and the molecular linkages between focal adhesions, myosin and F-actin, which are key mediators of contact guidance (*10, 49*), are likely to be important for extrusion into collagen matrices. Indeed, consistent with this hypothesis, increased matrix density and stiffness promote ECM alignment and mammary cell invasion into 3D matrices through a Focal Adhesion Kinase (FAK)-ERK signaling axis, where inhibition of FAK function by expression of dominant-negative FRNK reverts the invasive phenotype (*17*). Moreover, a recent study by Denardo and colleagues demonstrates that targeting FAK in PDA not only augments immunotherapy efficacy but also decreases the number of single PDA cells observed in the periductal space (*7*). Thus, this collective evidence prompted us to hypothesize that FAK signaling promotes cell extrusion and subsequent invasion associated with aligned ECM. To test the impact of FAK inhibition (FAKi) we first employed our *in vitro* epithelial-ECM interface platform. Consistent with our hypothesis, FAKi dramatically reduced cell extrusion, to the extent that very little single cell extrusion was observed at Day 1 in the FAKi group versus robust extrusion in the vehicle control group (Fig. 4g-i; see also comparison between Movies 8 and 9). This behavior was maintained through day 3, where extrusion and invasion was profoundly limited by FAK inhibition compared to controls, which exhibited remarkable extrusion and invasion (Supplementary Figure 4e), suggesting that FAKi indeed limits extrusion and subsequent invasion. However, we note that in pancreatic carcinoma cells lacking S1P_2_ receptor, FAK has also been suggested to regulate apoptosis associated with basal extrusions in monolayer culture (*34*). As such, we evaluated apoptosis levels in basally extruded cells to determine if our observed decreases in extrusion and invasion under FAKi could be due, either entirely or in part, to apoptosis. In the primary cells within 3D environments in this study, analysis of cleaved caspase-3 (CC3) revealed no differences in apoptosis between control and FAKi treated carcinoma cells (Supplementary Figure 4f). Interestingly, concomitant with the decrease in extrusion, we did not observe a reduction in organized collagen, especially TACS-3 like arrangements that were present around the epithelial structures in both control and FAKi groups (Fig. 4g,h), suggesting that even in the presence of TACS-3-like architectures, inhibition of FAK profoundly reduces the ability of cells to extrude and then invade along aligned ECM. We do however observe limited collective invasions in the FAKi group at later time points (Days 2-4) where single cell invasion remains severely impaired (Supplementary Fig. 4e), which is also consistent with growth and general motility-related quasi-ballistic invasion into aligned ECM resulting from cell crowding (*37*). Thus, our data suggests that FAKi cells still retain limited ability to invade from well-defined ductal structures but disrupting the key signaling node of focal adhesion force transmission and contact guidance by targeting FAK makes this process quite inefficient.

### FAK inhibition abrogates single cell extrusion and metastasis in PDA

Previous work using the *KPC* model demonstrated that FAKi leads to a decrease in fibrosis in PDA, with concomitant reduction in the number of invasive PDA cells observed in the periductal space (*7*). We therefore analyzed PDA FFPE sections from Vehicle vs. FAKi groups to interrogate cell extrusion and invasion in the context of collagen fiber organization to test our hypothesis that FAK influences early dissemination *in vivo*. Consistent with the general abrogation of fibrosis, FAK inhibition led to a decrease in fibrillar collagen in both early (1.5 month group) and end stage *KPC* mice (Fig. 5a-c) with a modest, yet significant, decrease in TACS-2 architectures and a modest, not significant, trend of less TACS-3 like architectures (Fig. 5d). These architecture are likely influenced by attenuation of FAK-dependent mechanotransduction in myofibroblasts that regulates the fibroinflammatory response in PDA (*50*). Thus, while collagen levels are reduced, TACS architectures are still present in the vast majority of ductal structures (Fig. 5c,d). In agreement with our hypothesis and *in vitro* data, there was a significant decrease in the frequency of single cell extrusions following inhibition of FAK (Fig. 5e), consistent with the observation of smoother and more organized ductal morphology in FAKi-treated mice (Fig. 5b), indicative of a less advanced or more limited progression of disease (Fig. 5a, b). Interestingly, the number of extrusions associated with both TACS-2 and TACS-3 showed decreasing trends following FAK inhibition. However, the difference was significant and much more pronounced for TACS-3 associated extrusions (Fig. 5f), suggesting that TACS-3-mediated extrusions and invasion may be more FAK-dependent. This is critical since collective data suggests that for robust dissemination of extruded cells invading through periductal TACS-2 architectures ultimately rely on TACS-3 architectures to serve as an exit point from the duct-adjacent space. Indeed, *KPC* mice undergoing FAKi treatment showed decreased metastasis to the liver ((*7*); Fig. 5g), consistent with the conclusion that FAK-dependent single cell extrusion and subsequent invasion through the stroma is a precursor to metastasis. Overall, these data implicate FAK as a key mechanotransduction node that influences the frequency of early single cell extrusion and subsequent invasion in PDA leading to enhanced metastatic dissemination. Thus, early dissemination and therefore early metastasis in PDA is driven, at least in part, by the local organization of periductal collagen and focal adhesion-dependent recognition of these ECM patterns by carcinoma cells.

**Fig. 5:**
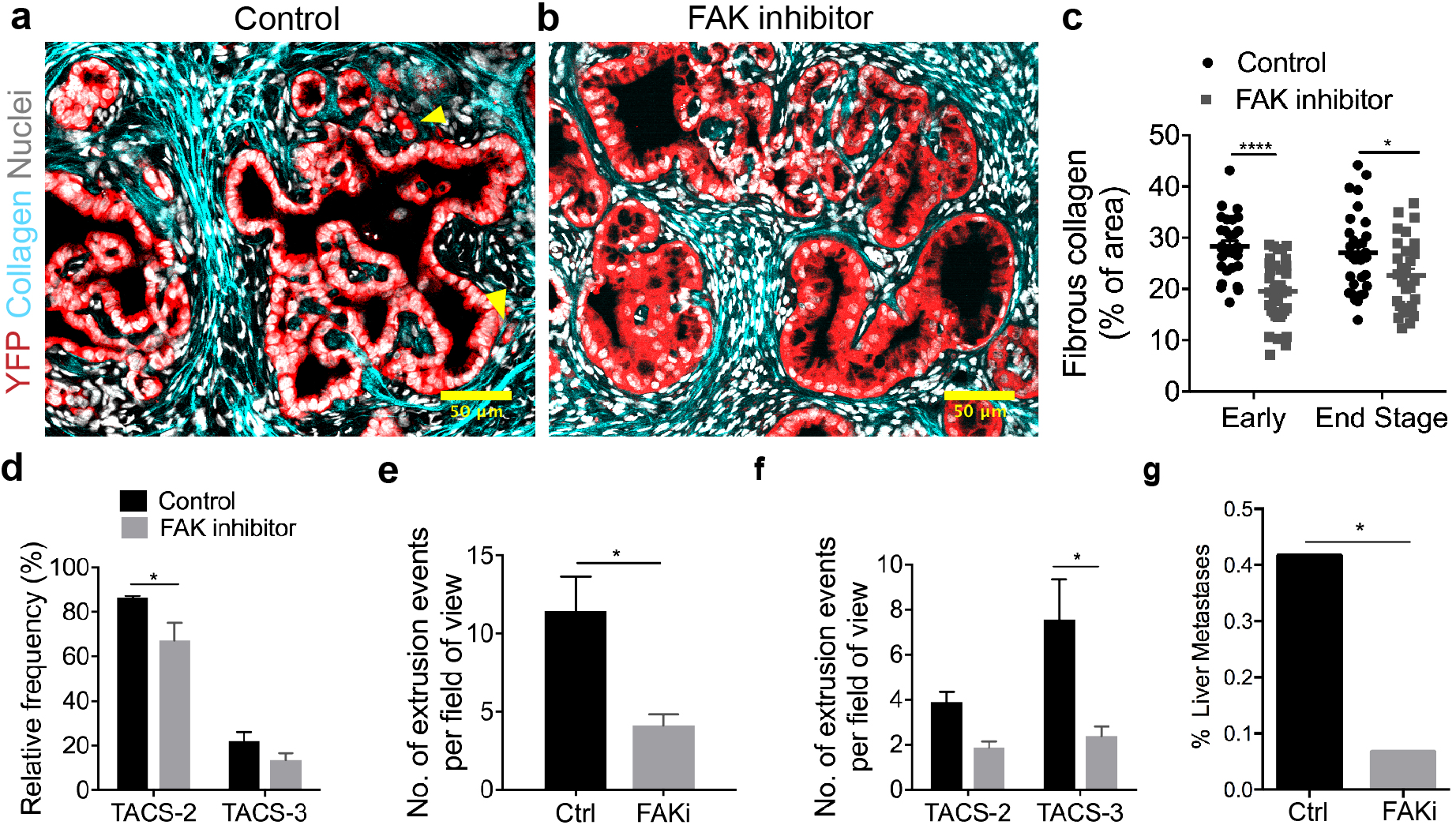
Inhibition of FAK alters frequency of collagen architectures, extrusion and metastasis in PDA: (**a,b**) Fluorescence micrographs of stained *KPCY* sections either treated with (a) vehicle control or (b) FAK inhibitor showing reduction in collagen deposition around ductal structures, smoother ductal boundaries and reduction in cell extrusion; *yellow arrowheads* indicate single cell extrusion events in the control sample. (**c,d**) Quantification of collagen content in *KPC* mice shows a reduction in deposition of fibrous collagen in both early (1.5 month) and end stage mice, with a (d) concomitant reduction in TACS-like architectures in *KPCY* mice. Note, the vast majority of PanIN lesions in FAKi treated mice still retain TACS2 and TACS3 architectures in spite of lower overall collagen levels. (**e**) Number of extrusion events quantified from Control and FAKi treated mice showing reduced extrusion by FAK inhibition. (**f**) Extrusion events for the control and FAK inhibited groups quantified as a function of TACS-2 or TACS-3-architectures showing a profound decrease in extrusion and into TACS3 regions. (**g**) Frequency of liver metastasis in control and FAKi treated *KPC* mice showing reduction in liver metastases with FAK inhibitor treatment (data are from *Jiang et. al* (*7*)). Data are mean +/− SEM (c-f), n=29-36 fields of view per group, ****p<0.0001, *p<0.05 by 2-way ANOVA and Sidak’s multiple comparison test for (c, d, f), n=3 per group, *p<0.05 for (e) by t-test, *p<0.05 by Fisher’s exact test for (g); Scale bars = 50 μm.

## Discussion

Our findings represent a key advancement in the understanding of how the desmoplastic ECM drives disease progression in PDA. We utilize combined MPE and SHG imaging to visualize and quantify collagen architectures in PDA and establish critical links between the periductal collagen organization and early invasion in PDA. Indeed, it is clear that the deposition and organization of periductal collagen plays an important role in the early dissemination and metastasis of PDA and may be utilized as a biomarker for disease progression; and one can envision using TACS to map and identify key features for tracking disease progression in other carcinomas as well (*51, 52*). Overall, these novel insights offer a bridge between the cell biology of basal epithelial extrusion, invasion through 3D environments, and clinical oncology and provide further impetus to the growing paradigm of stroma-targeted therapies to normalize cellular microenvironments in pancreatitis and cancer.

In this work, we identify and map previously established disease-relevant collagen signatures (i.e., TACS in breast tumors) in pancreatic disease. Both TACS-2 and TACS-3 manifest with largely straightened collagen fibers in PDA, albeit often with a looser organization for TACS-2 than was observed in mammary carcinomas. We note that collagen is highly anisotropic with a modulus that is orders of magnitude larger in the direction of alignment versus other directions (e.g. lateral deformations, bending, compression etc.) and widespread straightening of the collagen fibers has been associated with increased stiffness due to the well-established strain-stiffening behavior of fibrillar collagen (*53*). As such, elevated stiffness in itself has important implications in tumor progression (*17, 19, 54*), suggesting that the TACS architectures not only contribute to contact guidance but also likely promote disease progression through stiffness-regulated mechanotransduction signaling. Likewise, these straightened collagen networks are also consistent with increased tumor pressures and poor drug transport observed in PDA, and the previous descriptions that hyaluronan-driven swelling pressures are constrained by a stressed collagen network in PDA (*5, 55*). Furthermore, we observed cell extrusions in cancer-adjacent normal sections showing signs of inflammation and collagen deposition. One could posit that this acts to prime the adjacent regions for disease infiltration and simultaneous entrance into the metastatic cascade. Indeed, this development of the “ECM phenotype” may precede and even drive the “malignant phenotype” in these adjacent regions. Importantly, our findings suggest this is likely the case for fibrotic diseases like pancreatitis. Notably, the onset of induced pancreatitis appears to augment early dissemination in *KC* and *KPC* mice (*11*), and we found ample evidence of fibrotic collagen in the form of TACS-2 and TACS-3 in human pancreatitis, but with minimal extrusion (Fig. 2), suggesting that TACS alone may not be sufficient for robust cell extrusion in untransformed cells. Therefore, the risk factor posed by chronic pancreatitis for malignant transformation may at least, in part, be attributed to the generation of the “ECM phenotype” as a priming microenvironment for PDA to arise and disseminate from early stages, and that therapeutic approaches to modify the microenvironments in the pancreas could play a role in limiting early metastatic disease from PDA.

In terms of the cell-intrinsic pathways regulating the cell biology of basal extrusion from ductal epithelia, data suggest that sphingosine-1-phosphate (S1P) and Rho-mediated signaling play a role mediating actomyosin contractility during extrusion of epithelial cells in monolayer culture conditions (*34*). Likewise, loss of p120 catenin in *Kras* driven PDA also promotes epithelial cell extrusion with concomitant increases in fibrotic stroma in early disease in *KC^iMst1^* mice (*35*), which is consistent with our findings of stromal architecture guiding basal extrusions and invasion. Here we also show that in PDA, FAK is crucial for robust cell extrusion and subsequent invasion through organized ECM. Thus, basal extrusion is clearly a biophysical process requiring force imbalance, not dissimilar to the anisotropic forces driving contact guidance and cell-cell forces that can compete with cell-ECM forces (*10*). Taken together, our data suggests that this principle may be applied generally to study cell-ECM interactions and the balance of cell-cell and cell-ECM forces is a key factor driving extrusion, single and collective cell invasion and colonization. Along these lines, here, we introduced a microfluidics-based high-throughput assay that enables the study of delamination of carcinoma cells at the epithelial-collagen interface. This system provides a highly controllable platform to model the epithelial-ECM interface and can be used in the future to address fundamental developmental and disease biology and perform rapid drug screening. Certainly, it could be used to support additional studies to further establish and define focal adhesion-cytoskeleton-contractility related pathways necessary for fundamental extrusion and motility process and to determine the differential effects of various stromal components in this process, which are much more readily controllable *in vitro* than *in vivo.*

In terms of therapy, our findings provide further evidence to support the development of stroma-targeting therapies (STTs) to treat desmoplastic diseases like PDA (*4, 5, 7, 41*). Indeed, we and others have recently developed regimens to target the stroma in autochthonous established PDA (*5, 7, 8, 41*) to provide strong consensus evidence of the potential for disrupting the stroma. For instance, we recently demonstrated that Halofuginone can help normalize CAF behavior to robustly decrease the fibrotic response in PDA while also increasing anti-tumor immunity (*41*). Likewise, antifibrotic therapy with Losartan has been shown to decrease fibrous ECM and has gone through promising Phase II trials (*56*), and multiple early trials with FAK inhibitors are ongoing for PDA (e.g. NCT02758587, NCT04331041, NCT03727880). Thus, therapeutic strategies to re-engineer TMEs to move them toward normalization may have a role not only in limiting continued dissemination from unresectable invasive disease during treatment, but also in very early disease. As much needed approaches for early PDA detection advance, it may be necessary to couple stromal and molecular therapies (such as STT and FAKi approaches) for early treatment in order to limit early dissemination. Likewise, it may be beneficial to employ STT strategies to treat precursor diseases such as pancreatitis. We observed robust fibroinflammatory activity in pancreatitis with establishment of ECM that results in stromal architectures that are primed for robust dissemination prior to transformation such that they can facilitate disease spread from the earliest onset of disease. Removing these architectures in at risk patients may be a strategy to impede early dissemination in persons who later develop PDA. Indeed, stroma normalizing approaches could be explored as a prophylactic drug to dismantle the microenvironment that is likely to promote disease progression and early metastasis.

## Materials and Methods

### Human and mouse pancreatic tissues and tumors

Normal and diseased pancreatic biopsies (PDA and chronic pancreatitis) from human patients were obtained as formalin-fixed paraffin-embedded (FFPE) sections in the form of a tissue with associated pathology information (Staging, grading etc.; US Biomax, Inc). In addition, freshly resected PDA, CAN, and chronic pancreatitis sections from the clinic were obtained from BioNet (University of Minnesota) in compliance with approved Institutional Review Board protocols. Fresh tissues were kept on ice in cell culture media for transportation, formalin-fixed immediately upon arrival and subsequently paraffin-embedded and sectioned for analysis.

Genetically engineered *Kras^G12D/+^ ;Tp53^R172H/+^ ;Pdx1-Cre* (*KPC*) mice were used as a faithful mouse model for pancreas cancer that recapitulate key stromal dynamics and metastatic spread observed in human disease (*5, 24*). Pancreatic tumor samples were obtained from variants of the *KPC* model (*KPCT: KPC; ROSA26^LSL-tdTomato/+^ or KPCG*: *KPC; ROSA26^LSL-ZsGreen1/+^ or KPC; ROSA26^LSL-YFP^*) with fluorophores expressed specifically in pancreatic carcinoma cells. All animal studies were approved by the University of Minnesota Institutional Animal Care and Use Committee.

### Cell Culture

Primary PDA cell lines were derived from pancreatic tumors in *KPCG* or *KPCT* mice were derived as previously described (*5*). Following establishment and purification, cells where maintained in high glucose DMEM supplemented with 10% FBS. 3D culture was conducted as we described previously (*17, 27*). Briefly, these primary pancreatic carcinoma cells were embedded in 3mg/mL collagen-I (Corning) matrices or in Matrigel (Corning), diluted in complete medium to a concentration of 4mg/mL, for 2 days before imaging. (See the “Engineering microtissues to analyze cell extrusion and invasion” section for details on fabrication of collagen matrices).

### Staining and imaging of archival tissues

FFPE sections were imaged on a custom-built multi-photon laser scanning microscope with 4-channel detection with interchangeable 440/20, 460/50, 525/50, 525/70, 595/50, 605/70, and 690/50 filters, (Prairie Technologies / Bruker) and a Mai Tai Ti:Sapphire laser (Spectra-Physics) to simultaneously produce MPE and SHG to visualize cells and collagen, respectively, which has been described in detail previously (*57*). To capture SHG from collagen, samples were imaged at 880-900nm and harmonic signal captured with a blue bandpass filter with a 440 or 450nm center wavelength. For human samples, the overall architecture of the tissue was visible by imaging endogenous fluorescence and was captured with a green emission filter at the same wavelength of excitation of 880-900nm. Alternately, some samples were stained with FITC-conjugated anti-pan Cytokeratin antibody (1:100, Sigma) to visualize carcinoma cells.

Since ZsGreen1 retained its fluorescence post-fixation, FFPE sections obtained from *KPCG* mice were imaged without additional staining for the carcinoma cells. ZsGreen1 fluorescence was excited by 880-900nm 2-photon excitation and detected through the green emission filter. Sections from *KPCT* mice were additionally stained with an anti-RFP antibody (1:100, Abcam) and Alexa-conjugated secondary antibodies (Thermo Fisher) were used to visualize the tdTomato^+^ carcinoma cells post-fixation. The tdTomato fluorescence was imaged at 800nm two-photon excitation using the red emission filter sets (595/50). Note, all tdTomato or zsGreen fluorescence is pseudo-colored red in image throughout the manuscript for consistency of identifying carcinoma cells.

For staining FFPE sections (Cytokeratin or RFP), slides were first dewaxed and rehydrated in xylene and ethanol followed by antigen retrieval in Citrate buffer (pH 6.0) for 20mins in a steamer. Subsequently, the sections were blocked in 5% goat serum and incubated with the primary antibody overnight with 1% goat serum. The secondary was allowed to incubate for 40mins with 1:30000 bisbenzimide or Hoechst before washing and mounting with Prolong Gold. Alternatively, slides were subjected to antigen retrieval by steam heating in 1X Trilogy (Cell Marque). Slides were permeabilized with 0.025% Triton-X-100 (Roche) in 1X TBS (Bio-Rad) (TBS-T) and blocked for 1h at RT with 5% goat serum (Vector Labs) in 1X TBS-T. After blocking, for immunofluorescence, slides were incubated overnight in a humidified chamber at 4 °C in TBS-T with 1% Fatty acid free-BSA (Fisher Scientific) with anti-mouse/human antibodies. The following primary antibodies were used: 1:50 mouse anti-Cytokeratin Pan-FITC (MilliporeSigma, Human), 1:100 mouse anti-Pan-Keratin (Cell Signaling, Mouse), 1:200 rat anti-mouse CD31 (Dianova, Mouse), 1:400 rabbit anti-CD31/PECAM-1 (Novus Biologicals, Human), 1:400 rabbit anti-RFP (Abcam, Mouse), 1:200 rabbit anti-GFP (Life Technologies, Mouse). Subsequently, slides were washed and incubated for 1h at RT with 1:200 Alexa-fluor secondary antibodies: goat anti-rabbit 488 (Life Technologies), goat anti-rat 568 (Life Technologies) along with 1:500 Draq5 (Biolegend) nuclear stain followed by wash steps and mounting with Prolong Gold (Life Technologies).

### Quantification of collagen content and architecture

For unbiased quantification of collagen content and TACS, multiple SHG images were taken at random locations on several slides obtained from different *KPCT or KPCG* mice or human patients. These images of different regions represented various grades or stages of PDA progression such as normal, adjacent normal, PanIN, undifferentiated PDA or well-differentiated PDA, which were determined by combining pathological assessment (also in case of the human samples) and inspection of hematoxylin and eosin (H&E) stained serial sections of the same tumor. The SHG images were then run through a custom Matlab code that dynamically thresholds images to retain collagen positive pixels in a binary image. The percentage of positive pixels were then calculated and used as a measure of fibrous collagen content. Quantification of TACS-2 and TACS-3 was obtained using CurveAlign (LOCI), a freely available software that calculates the angular distribution of collagen fibers with respect to a user-defined ductal boundary (*58*). To obtain the frequencies of TACS-2 and TACS-3 occurrence, entire millimeter-scale regions of the human biopsies were imaged using MPE and SHG, reconstructed and the number of total ducts, the number of TACS-2 and TACS-3 positive ducts were noted and corresponding percentages obtained.

### Live cell imaging of tumor explants

Freshly excised tumor slices were obtained from *KPCT* and *KPCG* mice (note, all tdTomato or zsGreen fluorescence is pseudo-colored red throughout the manuscript for consistency of identifying carcinoma cells). Live cell imaging of tumor slices was performed similar to time-lapse MPE methods described previously (*27*). Briefly, the tumor was sliced into a few thin sections using a vibratome, set on a 35mm dish with a slice anchor (Warner Instruments) and overlaid with L-15 media supplemented with 10% fetal bovine serum (FBS), penicillin-streptomycin, plasmocin, fungizone and 10μg/ml soybean trypsin inhibitor (Sigma). Imaging was subsequently performed on the multiphoton microscopy setup described above with a custom-built temperature-controlled stage insert (*59*) for 6-12h with a time interval of 20mins between frames at 880-900nm excitation wavelength and emission captured in the green (*KPCG*) or red (*KPCT*) and the blue (SHG) channel. Analysis and 3D rendering of Z-stacks, visualization of time-lapse imaging data and image processing were done in Fiji. For live imaging, Z-stacks were generally obtained at a step size of 4-5μm, covering a total depth of about 80-100μm.

### Analysis of cell extrusion *in vivo*

For analysis of cell extrusion, unbiased images of CAN, PanIN and differentiated PDA regions (17-54 fields of view/group) were obtained from pancreatic tumor sections (6-13/group) from different *KPCT* or *KPCG* mice (7-9 mice/group). Within each field of view (300μmX300μm or 600μmX600μm, the number of ductal structures, the percentage with organized periductal collagen (either TACS-2 or TACS-3) and the total number of extrusion events were calculated. An extrusion event was defined as single (or dividing) disseminated cell in the periductal stroma (fully extruded) or a single or pair of cells protruding out from the smooth boundary of a duct (partially extruded). The association of partially extruded cells with TACS-2 or TACS-3 positive areas or that of fully extruded cells with aligned collagen in the stroma was determined by visual inspection, keeping in mind quantitative measures described in Fig.1. Morphometric analysis of disseminated cells was performed manually in Fiji, including only single cells fully extruded into the stroma for the analysis.

### Engineering microtissues to analyze cell extrusion and invasion

Matrigel microtissues approximately 150-200μm in diameter were fabricated modifying previously established protocols (*60, 61*). Matrigel (Corning) was thawed on ice overnight and diluted to a concentration of 6 mg/mL with Dulbecco's phosphate-buffered saline (DPBS). At 4°C, the Matrigel solution was partitioned into droplets using a flow-focusing polydimethylsiloxane (PDMS) (Dow Corning) microfluidic device. The continuous phase from the droplet generation (FC-40 with 2% 008-FluoroSurfactant, Ran Biotechnologies), was collected with the droplets in a low retention Eppendorf tube and polymerized for 30 minutes at 37°C. The oil phase was removed and the Matrigel droplets were resuspended in 1x DPBS with a manual micropipette.

Droplets were then moved to agarose microwells, fabricated following previously established protocols (*61*). Briefly, polystyrene multi-well plates were coated with 2% agarose and dehydrated in a sterile laminar flow hood overnight. PDMS stamps with 300 μm diameter posts were plasma treated for 2 minutes to produce a hydrophilic surface and sterilized with boiling water. Agarose solution was pipetted into each well and the hydrophilic PDMS stamp placed immediately onto the molten agarose. After cooling for 5 minutes, stamps were removed gently from the polymerized agarose and hydrated with 1 x DPBS. Wells were washed with appropriate culturing media prior to adding microtissues with a manual micropipette.

Once inside the microwells, droplets were coated with primary *KPCT* or *KPCG* cells at a concentration of 100,000 cells/well. These microtissues were cultured in complete DMEM for 7 days to allow cells to adhere to the outside of the droplets and form luminal structures. Droplets were then treated daily with either 1μM FAK inhibitor VS-4718, or with vehicle control (DMSO) for three days, after which they were embedded in 4mg/mL 3D collagen-I gels (Corning), with continued inhibitor or DMSO treatment. Briefly, collagen gels were generated by mixing collagen-1 with an equal volume of 100mM HEPES buffer in 2X DPBS, as previously described (*27, 37, 59*). Droplets were added to this neutralized solution, which was then aliquoted out in 475uL increments into a 12-well culture plate and allowed to polymerize for 30 minutes at room temperature. Constructs were moved to 37°C for 4 hours before overlaying media. Embedded droplets were imaged using MPE and SHG imaging over days 0-4 days post-embedding. Invasion events were quantified by counting the number of cells protruding from luminal structures. Laminin staining was performed with 1:100 anti-laminin Ab (ThermoFisher) and 1:200 Alexa-fluor secondary antibody as described in the “Staining and imaging of archival tissues” section.

### FAK inhibition in *KPC* mice

Control and FAKi samples have been previously described (*7*). Briefly, 50mg/kg VS-4718 was administered by oral gavage twice a day in a formulation with 0.5% carboxymethyl cellulose and 0.1% Tween-80 (Sigma) in sterile inhibitor.

## Statistical analysis

Multiple groups were compared by ANOVA, followed by the Tukey posthoc analysis, or the non-parametric Kruskal-Wallis test with Dunn’s post-hoc testing, as dictated by the size and distribution of the data. Similarly, 2-way ANOVA with Sidak’s multiple comparison test was used to compare multiple groups across different conditions. The non-parametric Mann Whitney test was employed for testing null hypotheses between two groups for small datasets. Number of data points for each experiment, the specific statistical tests, and significance levels are noted in the figure text.

## Acknowledgments

PPP and this work was supported by Research Scholar Grant, RSG-14-171-01-CSM from the American Cancer Society and the NIH (R01CA245550 to DKW and PPP, R01CA181385 to PPP and U54CA210190 University of Minnesota Physical Sciences in Oncology Center, Project 2 to PPP and Core 1 to DKW). This work was also supported by the Randy Shaver Research and Community Fund (PPP), Masonic Cancer Center (PPP), a University of Minnesota Doctoral Dissertation Fellowship (AR), and grants from the UMN Institute for Engineering in Medicine (PPP and DKW). The content of this work is solely the responsibility of the authors and does not necessarily represent the official views of the NIH or other funding agencies. The authors thank members of the Provenzano laboratory for insightful comments regarding this work.

## Author Contributions

PPP initially conceptualized the study. AR, MKC, and PP participated in the design of the study. AR, MKC, NJR, HRR, ALC, HJ, DGD, DKW, and PPP participated in the design of experiments. AR, MKC, NJR, ALC MC, KBE, EAE, JHS, NRR, and HJ generated unique reagents, performed experiments and/or analysis. AR developed quantitative metrics and algorithms. ALC and DKW developed the microfluidic platform and developed its use here with AR, MKC, and PPP. JH and DGD generated murine tissues from FAK inhibition studies and assisted in experimental design and data interpretation. PPP and DGD secured funding. AR, MKC, and PPP wrote the manuscript, with significant edits by NJR. All authors read and contributed comments to the final manuscript. PPP oversaw all aspects of the study.

## Movie Legends

**Movie 1:** Surface rendering from a 3D multiphoton fluorescence micrograph of tdTomato expressing pancreatic carcinoma cells (*red*) from a live *KPCT* tumor explant showing extruded single cells from ductal structures.

**Movie 2**: Combined MPE and SHG imaging a live *KPCT* tumor explant showing fluorescent carcinoma cells (*red*) in ductal structures surrounded by collagen (*cyan*) undergoing coherent angular motion.

**Movie 3**: Combined MPE and imaging a live *KPCG* tumor explant showing two extruded carcinoma cells of small size and aspect ratio (*red*) interacting with periductal collagen (*cyan*) and switching phenotype.

**Movie 4**: Combined MPE and imaging a live *KPCT* tumor explant showing elongated single carcinoma cells (*red*) orienting and migrating along aligned periductal collagen tracks (*cyan*).

**Movie 5:** Combined MPE and imaging a live *KPCT* tumor explant simultaneously showing partially and fully extruded single carcinoma cells (*red*) associated with periductal collagen (*cyan*).

**Movie 6:** Combined MPE and SHG of *KPCT* microtissue embedded in a collagen matrix (*cyan*) at day 0 showing a stable epithelial morphology for carcinoma cells (*red*). Movie is imaging in 20min intervals over 12 hours.

**Movie 7:** Combined MPE and SHG of *KPCT* microtissue embedded in a collagen matrix (*cyan*) at day 3 showing extrusion and subsequent invasion of carcinoma cells (*red*). Movie is imaging in 20min intervals over 8 hours.

**Movie 8:** Combined MPE and SHG of *KPCT* microtissue embedded in a collagen matrix (*cyan*) under control conditions showing extrusion and subsequent invasion of carcinoma cells (*red*). Movie is imaging in 10min intervals over 10 hours.

**Movie 9:** Combined MPE and SHG of *KPCT* microtissue embedded in a collagen matrix (*cyan*) under FAK inhibition (FAKi) conditions showing inhibition of extrusion and invasion of carcinoma cells (*red*). Movie is imaging in 10min intervals over 10 hours.

## Supplementary Information

**Supplementary Fig. 1:**
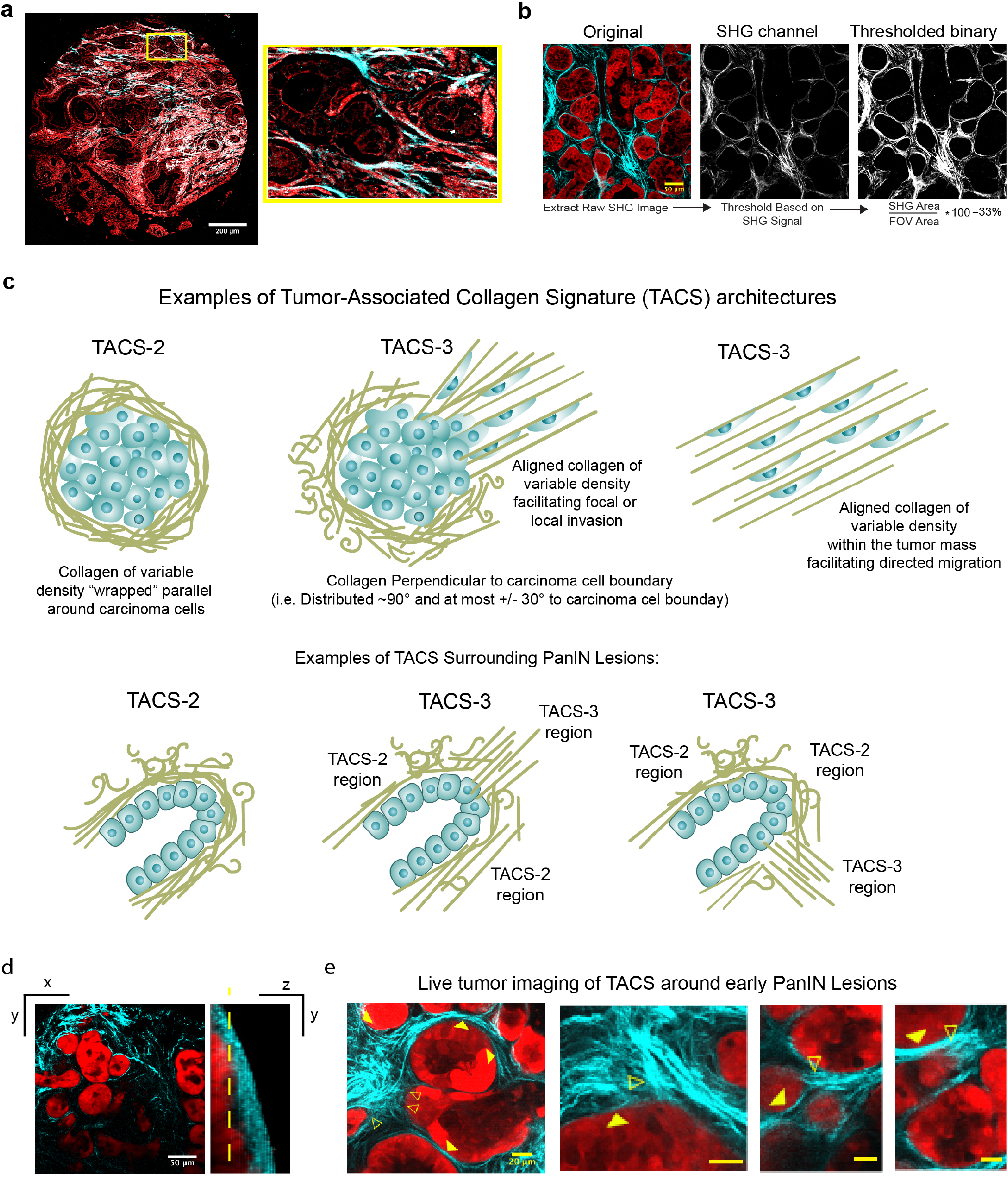
Imaging and analysis of collagen patterns in human and murine PDA: (**a**) Imaging and quantification of whole human biopsy samples (from TMAs with associated pathology staging and grading) for quantification of collagen signatures. Yellow box shows magnified view of the boxed region showing fibrous collagen patterns around ductal structures. (**b**) Example of fibrous collagen quantification from a *KPCT* sample. (**c**) Examples of Tumor-Associated Collagen Signatures (TACS) architectures and examples of these architectures near PanIN lesions. (**d**) 3D imaging of a PanIN section of a *KPCT* mouse with xy and yz views showing collagen wrapped around ductal structures-yellow dashed line shows the z plane of the xy image (**e**) Live multiphoton excitation of tdTomato expressed in carcinoma cells (*red*) with combined second harmonic generation of collagen (cyan) showing four additional examples of regions of the duct are positive for TACS-2 (solid arrowheads) while adjacent regions of the ducts are positive for TACS-3 (open arrowheads), demonstrating that collagen surrounding “pre-invasive” PanIN lesions possess ECM patterns that are known to promote invasion of pancreatic carcinoma cells. Scale bars = 200 μm (a), 50 μm (b, d), and 20 μm (e).

**Supplementary Fig. 2:**
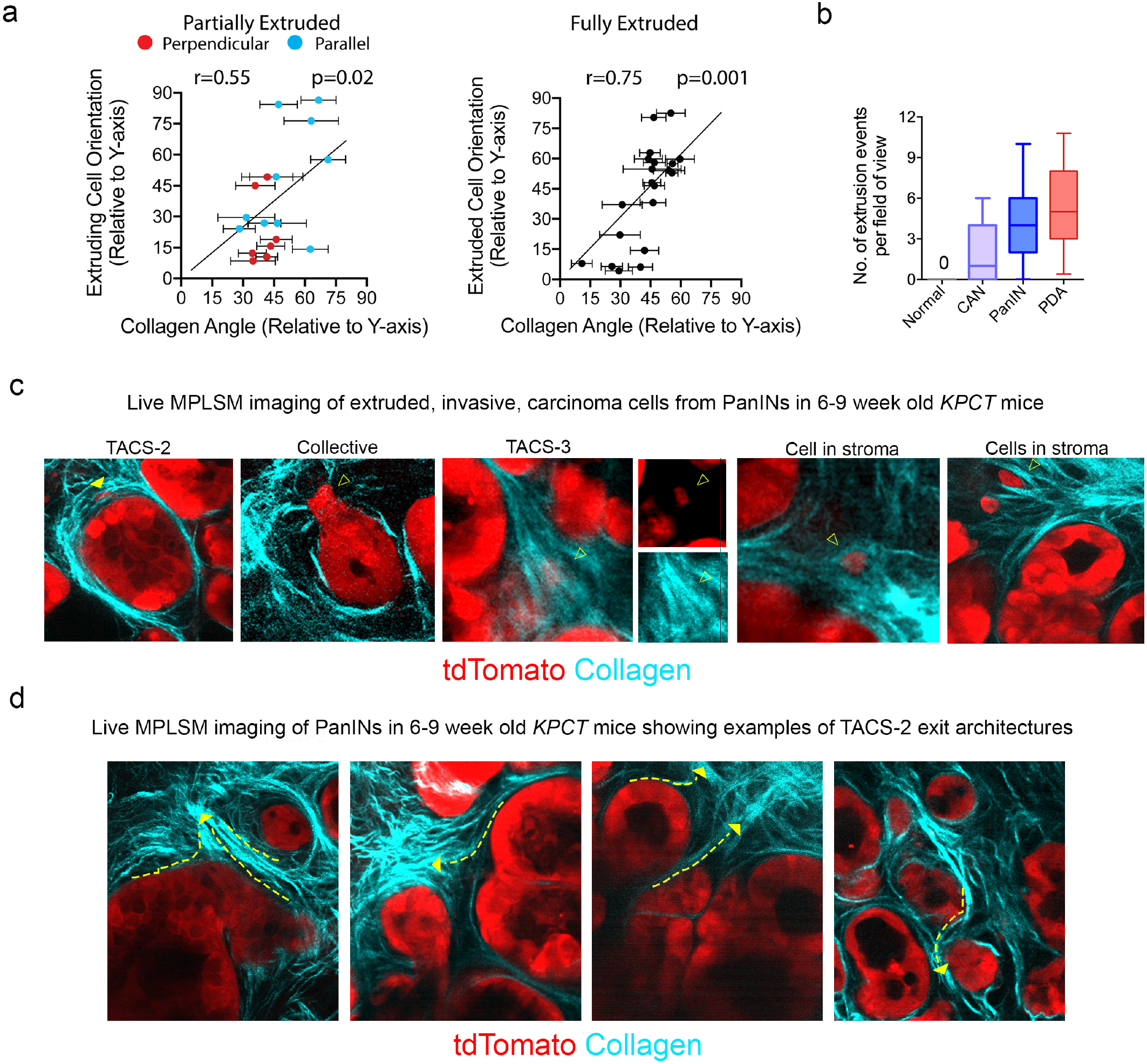
Extruded and invasive cells associated with TACS architectures: (**a**) Correlation between partial or extruded cells and collagen orientation showing associations between cell aspect ratio direction and fiber alignment. (**b**) Quantification of extrusion events for normal, CAN, PanIN and well-differentiated PDA from *KPCT* and *KPCG* mice. (**c**) Live tissue multiphoton laser-scanning microscopy (MPLSM, i.e multiphoton excitation of tdTomato expressed in carcinoma cells (*red*) with combined second harmonic generation of collagen (*cyan*)) in young mice demonstrating extrusion events associated with aligned collagen regions. Again, we note that regions of the duct are positive for TACS-2 while adjacent regions of the ducts are positive for TACS-3. (**d**) Live MPE and SHG imaging identifying TACS3 “exit points” from TACS2 regions. Arrows indicate examples of TACS2 to TACS3 paths. Scale bars = 20 μm.

**Supplementary Fig. 3:**
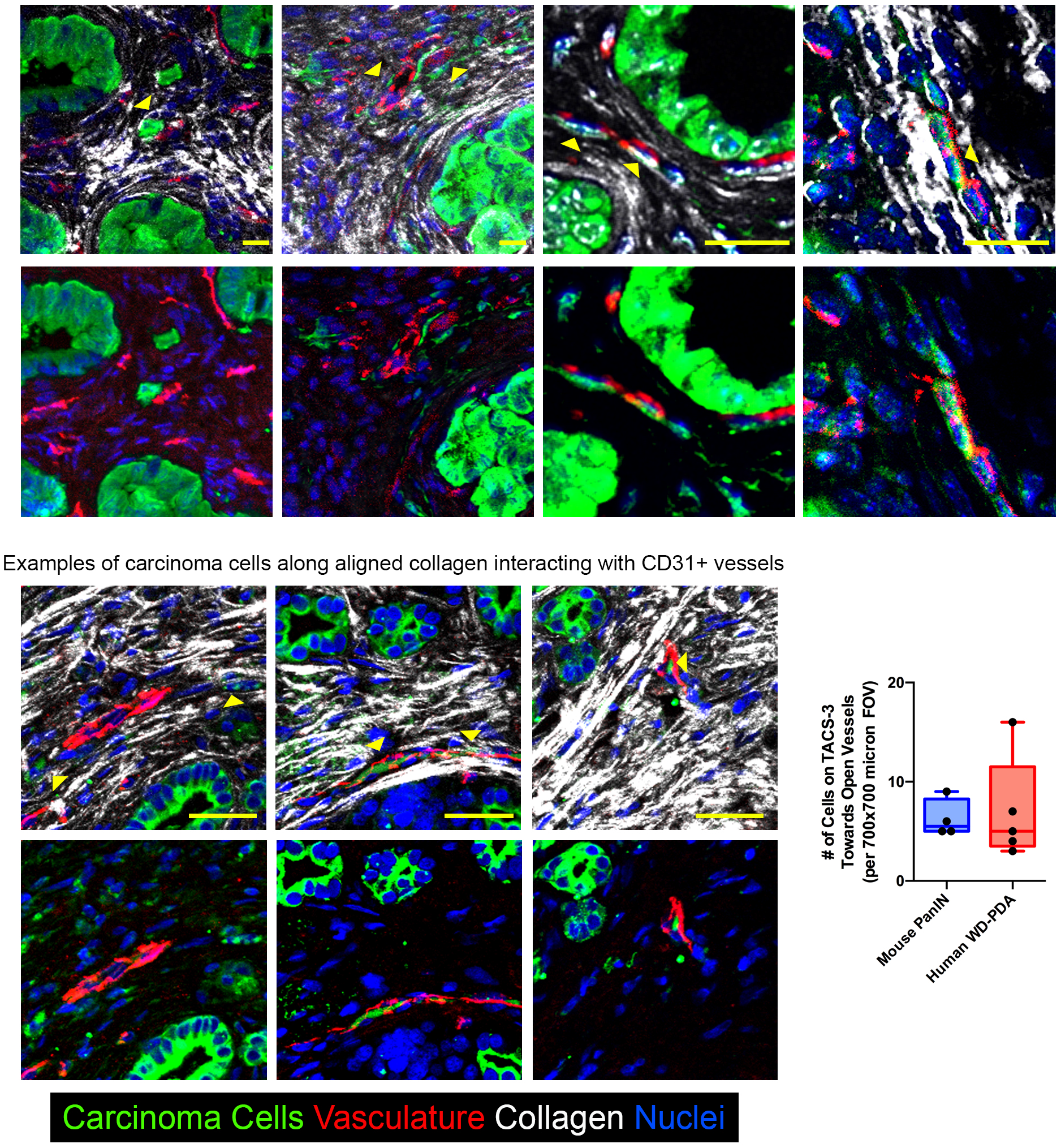
Extruded cell invading along aligned collagen to blood vessels. (***Top***) Immunofluorescence micrographs of *KPCT* tumor sections stained with RFP (*shown in green*), CD31 (*red*), and DRAQ5 (*blue*) demonstrating single extruded cells interacting with aligned periductal collagen (*white*) directed to blood vessels associated with early stage PanIN lesions; *Second row* shows immunofluorescence staining without collagen SHG signal. (***Bottom***) Immunofluorescence micrographs of well-differentiated human PDA stained with Cytokeratin (green), CD31 (*red*), and DRAQ5 (blue) demonstrating aligned extruded cells following collagen tracks (*white*) leading to blood vessels; *Fourth row* shows immunofluorescence staining without collagen SHG signal.. Scale bars = 20 μm (*top panels*) and 50 μm (*bottom panels*).

**Supplementary Fig. 4:**
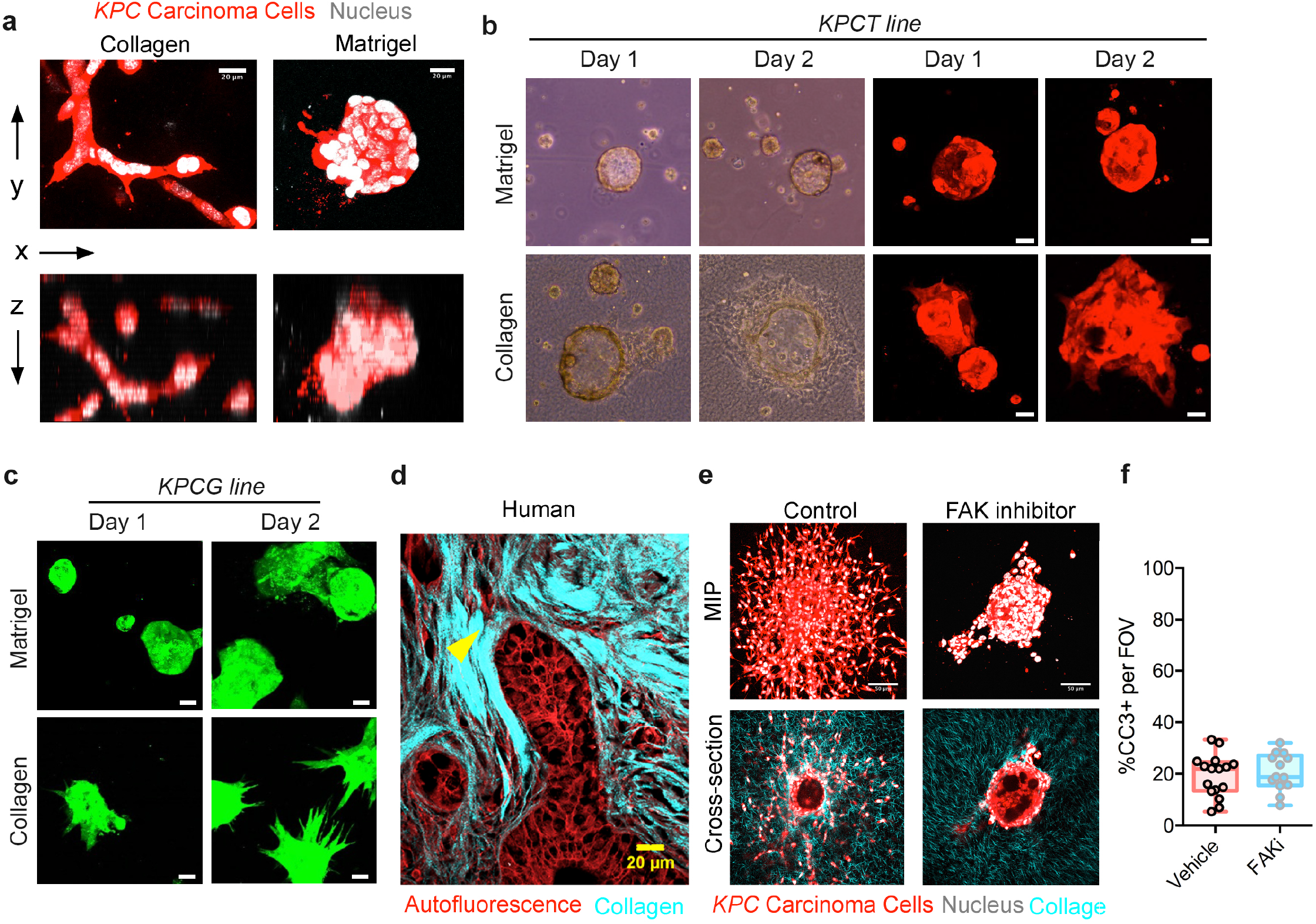
Single cell extrusion and dissemination *in vitro* and *in vivo*: (**a**) MPE micrographs of primary *KPCG* cells (pseudo-colored red) cultured in 3D collagen and 3D Matrigel showing contrasting morphologies with cells forming tight ductal-like clusters in Matrigel in contrast to more tubular, spindle-shaped, protrusive structures in 3D fibrous collagen matrices. (**b**) *KPCT* microtissues formed with microfluidic technology embedded in either Matrigel or collagen matrices showing that in predominantly basement membrane environments (i.e. Matrigel) the duct-like microtissues largely retain their epithelial phenotype while in collagen they begin to expand and invade. (**c**) Microtissues formed using a more invasive primary line (*from a KPCG mouse)* again shows a robustly invasive phenotype in collagen matrices while in Matrigel the microtissues retain a rounded non-invasive morphology. (**d**) A section from a human PDA sample shows collective invasion of the ductal boundary and subsequent single cell extrusion along TACS-3 (yellow arrowhead), akin to the group protrusions and single cell invasion observed in the epithelial organoids. (**e**) Combined MPE+SHG imaging presented as maximum intensity projection and cross-sectional views of cell extrusion and invasion from epithelial organoids into the surrounding collagen in Control and FAK inhibitor treated conditions at Day 3, demonstrating the drastic differences in extrusion efficiency. (**f**) Evaluation of apoptosis from cleaved caspase-3 (CC3) immunofluorescence showing no differences between control conditions and FAK inhibitor (FAKi) treatment. Scale bars= 20 μm (a, d) and 50 μm (b, c and e).

